# Plaque microbial community differences in rampant dental caries after exclusion of a reference-supported *Streptococcus mutans* ASV: an analysis of public 16S rRNA sequencing data

**DOI:** 10.64898/2026.09.02.748957

**Authors:** Zhiyi Zhuo, Xiaoguang Xu

## Abstract

**Background:** Dental caries is increasingly understood as an ecological biofilm disorder rather than solely the consequence of a single organism. Although *Streptococcus mutans* is strongly implicated in cariogenesis, it remains unclear whether caries-associated plaque-community differences persist beyond this organism.

**Methods:** We reanalysed publicly available supragingival plaque 16S rRNA sequencing data from 88 preschool children: 44 with rampant caries and 44 who were caries-free (BioProject PRJNA1141721). Single-end DADA2 processing yielded 25,669 amplicon sequence variants (ASVs), including one ASV identified as *S. mutans* by exact matching within the sequenced region to eHOMD RefSeq v15.23. The pre-specified primary analysis compared Bray-Curtis community composition after exclusion of this ASV.

**Results:** Non-*S. mutans* community composition differed between groups (PERMANOVA pseudo-R^2^=0.0462, pseudo-F=4.16, P<0.001), without evidence of unequal multivariate dispersion (P=0.796); rarefaction produced a nearly identical result. Shannon diversity did not differ (P=0.486). The reference-supported *S. mutans* ASV was detected in 8/44 rampant-caries and 42/44 caries-free samples and had lower relative abundance in rampant caries. Among 758 prevalence-filtered non-*S. mutans* ASVs, 239 non-structural-zero ASVs were FDR-significant, with more showing lower than higher bias-corrected abundance in rampant caries (176 versus 63); 239 additional ASVs were classified as structural zeros. Individual ASV findings were sensitive to prevalence filtering. Adjustment for *S. mutans* abundance attenuated the global caries-group association (marginal pseudo-R^2^=0.0149, P=0.085) in the presence of strong collinearity.

**Conclusions:** Rampant caries was associated with a modest plaque-community compositional difference after exclusion of a reference-supported *S. mutans* ASV; the result was robust to rarefaction and was not accompanied by a corresponding difference in Shannon diversity. These findings support a community-level ecological interpretation but do not establish statistical independence from *S. mutans* or a causal role for individual taxa.

## Introduction

Dental caries is a biofilm-mediated disease in which frequent fermentable-carbohydrate exposure, acid production, and ecological selection can move plaque toward sustained demineralization. This ecological framing emphasizes the collective properties of the plaque community, its local environment, and its functional responses rather than assigning disease to a single microbial agent [1-4]. Community composition may therefore differ across clinical states even when a summary measure of within-sample diversity does not differ.

*Streptococcus mutans* nonetheless has a central and well-supported role in cariogenic biofilm biology. Its carbohydrate metabolism, acid tolerance, and contribution to extracellular-matrix formation provide a mechanistic basis for its long-standing importance in caries research [2,5-7]. The continued relevance of mutans streptococci does not require that they be the only organisms associated with disease in every cohort, sample type, or 16S sequence window.

Studies based on culture-independent community profiling have identified broader plaque patterns associated with caries and have shown that caries may occur with low or undetectable *S. mutans* in some samples [3,8-11]. These observations are compatible with an ecological account of disease: organisms associated with clinical caries can vary while the relevant community context remains important. They do not, however, show that any observed community pattern is independent of *S. mutans* abundance or that particular taxa cause caries.

PRJNA1141721 provides public 16S rRNA sequencing reads from supragingival plaque of 88 preschool children, including 44 with rampant caries and 44 who were caries-free [12]. The original matched case-control study described group differences in oral microbiota. Reprocessing the raw archive offers an opportunity to address a distinct question with the handling of non-overlapping paired-end reads and taxonomic assignment.

Removal of a taxonomically supported ASV from a sequencing feature table does not represent biological removal of the organism and cannot establish statistical independence from its abundance. It does, however, provide a direct way to test whether the remaining plaque-community composition continues to differ between clinical groups.

The question is clinically and biologically relevant because plausible plaque-community differences may coexist with a well-established role for *S. mutans*. A remaining compositional difference could reflect correlated community structure, whereas failure to detect such a difference would not disprove the organism’s importance. The analysis therefore uses a focused ecological endpoint and reports *S. mutans*-specific analyses alongside it. Interpretation is confined to association in the public cohort and does not rely on unmeasured functional, clinical, or absolute-abundance mechanisms.

Importantly, the study is not designed to identify a universal caries microbiome. Oral biofilms are spatially organized, and caries-associated communities can differ by tooth surface, lesion status, population, and laboratory methods. The value of a transparent reanalysis is therefore not to impose a single signature on all settings, but to test a precisely stated question within one accessible cohort.

This perspective also aligns the language of the results with what 16S data can support: relative community patterns and taxonomic associations, rather than direct evidence of metabolic activity or clinical causation. It also places the present result alongside, rather than above, complementary culture-based, molecular, and longitudinal designs.

We therefore reanalysed the publicly available plaque sequencing data to test whether a caries-associated community-composition signal persisted after exclusion of a reference-supported *S. mutans* amplicon sequence variant (ASV). The pre-specified primary endpoint was the difference in Bray-Curtis composition of the non-*S. mutans* ASV community between rampant-caries and caries-free children. Secondary analyses considered alpha diversity, whole-community composition, the reference-supported *S. mutans* ASV, differential abundance, and sensitivity to analytical choices.

## Methods

### Study design, public data source and ethics

This secondary reanalysis used public sequencing data from BioProject PRJNA1141721. The original matched case-control study enrolled 88 preschool children (44 with rampant caries and 44 who were caries-free) and analysed supragingival dental plaque [12]. The original study reported matching of the case and control groups by age and sex, but individual age/sex values and matching identifiers were unavailable in the public metadata; the original matching could therefore not be incorporated into the reanalysis. The original publication states that all procedures were approved by the local Ethics Committee of the Affiliated Stomatological Hospital of Kunming Medical University (approval no. KYKQ2021MEC089) and that the children’s parents or legal guardians provided written informed consent [12]. No recruitment, intervention, or new data collection occurred for this reanalysis.

### Sequence retrieval and single-end processing

The public archive contained 10,394,430 paired-end read pairs. The first and second sequencing reads (Read 1 and Read 2) were 227 and 224 bp, respectively. Because their combined length was shorter than the expected approximately 461-bp V3-V4 amplicon, there was insufficient overlap for reliable paired-end merging. Read 2 was selected for single-end analysis before case-control comparisons based on slightly higher non-chimeric read retention and oral-reference exact-match performance; both reads supported the same unique *S. mutans* reference assignment. Expected primer sequences were not detected at the read starts, consistent with primer removal before public deposition. Read 2 sequences were processed in R 4.2.2 using DADA2 v1.26.0 [13]. Reads were filtered using maxN=0, maxEE=2 and truncQ=2, without fixed-length truncation or 5-prime trimming; PhiX reads were removed. Error models were learned from 500,000 bases before dereplication and ASV inference with DADA2. Chimeric sequences were subsequently removed using the consensus method. This produced 8,275,024 final non-chimeric reads and 25,669 ASVs.

### Taxonomic assignment and identification of S. mutans

Taxonomy was assigned with RDP Trainset 18. ASV sequences were compared with the corresponding sequenced Read 2 region of eHOMD RefSeq v15.23, an oral-site-focused reference resource [14,15]. One ASV exactly matched the *S. mutans* reference sequence within this region without an equally matching alternative oral species in the reference set; this ASV was therefore assigned to *S. mutans* for the primary analysis. Read 2 sequences were reverse-complemented only for broad RDP taxonomic assignment; comparisons with eHOMD used their native orientation. This assignment is limited to the sequenced Read 2 region and does not confer species-level resolution on the remaining single-end ASVs.

### Pre-specified community analyses

The statistical analysis plan was finalized before caries-group results were examined and was not externally preregistered. A random seed of 20260902 was used for permutation and resampling analyses. Analyses were performed for the whole ASV community and for the non-*S. mutans* community. For the latter, the reference-supported *S. mutans* ASV was removed from raw counts, the sample denominator was recalculated, remaining counts were converted to within-sample relative abundance, and Bray-Curtis distances were calculated from that re-normalized composition. The primary endpoint was the group term in non-*S. mutans* Bray-Curtis PERMANOVA with 9,999 permutations [16,17]. Multivariate dispersion was assessed with betadisper. Shannon diversity was compared by Wilcoxon rank-sum testing with rank-biserial effects and bootstrap confidence intervals. A single rarefaction sensitivity analysis used 49,000 reads per sample.

### *S. mutans*-specific and adjusted community analyses

Detection of the reference-supported *S. mutans* ASV was compared by Fisher’s exact test and relative abundance by Wilcoxon testing. A secondary marginal PERMANOVA included log10(*S. mutans* relative abundance + 10^-6). This adjusted analysis was interpreted cautiously because group status and *S. mutans* abundance were strongly correlated; it does not estimate an effect statistically independent of *S. mutans* abundance.

### Differential-abundance analysis

Pre-specified differential abundance included non-*S. mutans* ASVs detected in at least 9 of 88 samples (10.2%). Differential abundance was analysed with ANCOM-BC v1.4.0 [18], using the caries-free group as the reference category. Positive coefficients indicate higher bias-corrected abundance in rampant caries and negative coefficients indicate lower bias-corrected abundance in rampant caries. Benjamini-Hochberg q<0.05 defined FDR significance among non-structural-zero ASVs. Structural-zero ASVs were reported separately.

### Post hoc robustness analyses

The analysis using a prevalence threshold of at least 18 of 88 samples (20.5%), the *S. mutans*-adjusted differential-abundance model, and the eHOMD reference comparison of the 20 leading ASVs were post hoc. They were used to describe directional robustness and taxonomic support, not to replace the pre-specified analysis.

## Results

### Public dataset and sequence processing

All 88 public supragingival plaque samples were retained (44 rampant caries and 44 caries-free). Single-end processing of Read 2 yielded 8,275,024 final non-chimeric reads and 25,669 ASVs. One ASV was assigned to *S. mutans* with sequence-region-specific eHOMD reference support (Table 1).

**Table 1.** Study and dataset characteristics.

| Characteristic | Value | Context |
| --- | --- | --- |
| Participants | 88 | Original study / present reanalysis |
| Groups | 44 rampant caries; 44 caries-free | Original study |
| Sample | Supragingival dental plaque | Original study |
| Sequencing target | 16S V3-V4 | Original study |
| Public accession | PRJNA1141721 | Public archive |
| Sequencing read analysed | Read 2 (single-end) | Present reanalysis |
| Final non-chimeric reads | 8,275,024 | Present reanalysis |
| Final ASVs | 25,669 | Present reanalysis |
| Reference-supported *S. mutans* ASV | 1 | Present reanalysis |
| Individual age/sex | Not publicly available for reanalysis | Limitation |

### Plaque community composition differed by caries status after exclusion of the reference-supported *S. mutans* ASV

The pre-specified non-*S. mutans* Bray-Curtis PERMANOVA showed a difference in community composition between groups (pseudo-F=4.16, pseudo-R^2^=0.0462, P<0.001; Table 2). Mean distance to group centroid was 0.426 in the rampant-caries group and 0.430 in the caries-free group (P=0.796), providing no evidence of unequal multivariate dispersion. The rarefaction sensitivity analysis was nearly identical (pseudo-R^2^=0.0461, P<0.001). Whole-community composition was also associated with group (pseudo-R^2^=0.0489, P<0.001).

**Table 2.** Core ecological results.

| Analysis | Feature set | Estimate/test | 95% CI | P | Role |
| --- | --- | --- | --- | --- | --- |
| Community composition | Non-*S. mutans* | PERMANOVA: pseudo-$R^2=0.0462$; pseudo-$F=4.16$ | - | <0.001 | Primary |
| Multivariate dispersion | Non-*S. mutans* | Mean centroid distance: 0.426 vs 0.430 | - | 0.796 | Diagnostic |
| Rarefaction sensitivity | Non-*S. mutans* | PERMANOVA: pseudo-$R^2=0.0461$ | - | <0.001 | Sensitivity |
| Shannon diversity | Non-*S. mutans* | Rank-biserial: 0.087 | -0.162 to 0.335 | 0.486 | Secondary |
| Community composition | Whole community | PERMANOVA: pseudo-$R^2=0.0489$; pseudo-$F=4.42$ | - | <0.001 | Secondary |
| Shannon diversity | Whole community | Rank-biserial: 0.089 | -0.160 to 0.336 | 0.476 | Secondary |
| Adjusted community composition | Non-*S. mutans* | Marginal PERMANOVA: pseudo-$R^2=0.0149$ | - | 0.085 | Secondary |
| *S. mutans* detection | Reference-supported ASV | Fisher OR: 0.0116 | 0.0011 to 0.0583 | $2.90 \times 10^{-14}$ | Secondary |
| *S. mutans* abundance | Reference-supported ASV | Rank-biserial: -0.940 | - | $2.47 \times 10^{-15}$ | Secondary |

### Shannon diversity did not differ between groups

Non-*S. mutans* Shannon diversity did not materially differ (median 4.30 in rampant caries versus 4.25 in caries-free samples; rank-biserial=0.087, 95% CI -0.162 to 0.335; P=0.486). Whole-community Shannon diversity was likewise not different (P=0.476).

### The *S. mutans* ASV showed an inverse association with rampant caries

The reference-supported *S. mutans* ASV was detected in 8/44 rampant-caries and 42/44 caries-free samples (odds ratio=0.0116, 95% CI 0.0011-0.0583, P<0.001) and had lower relative abundance in rampant caries (P<0.001). In the pre-specified marginal PERMANOVA including measured *S. mutans* abundance, the caries-group term was attenuated (marginal pseudo-R^2^=0.0149, P=0.085), while the *S. mutans* abundance term was similarly non-significant (marginal pseudo-R^2^=0.0151, P=0.082). Caries status and *S. mutans* abundance were strongly correlated (point-biserial r=-0.869; R^2^=0.755; VIF=4.09), limiting interpretation of conditional effects. The inverse *S. mutans* pattern is specific to this cohort and to the sequenced 16S region/reference assignment and is not interpreted as protective.

### Differential abundance was widespread among non-S. mutans ASVs

Under the pre-specified prevalence filter of at least 9 of 88 samples, 758 non-*S. mutans* ASVs were tested. Of the 758 tested non-*S. mutans* ASVs, 239 non-structural-zero ASVs were differentially abundant at FDR q<0.05: 63 had higher and 176 had lower bias-corrected abundance in rampant caries. An additional 239 ASVs were classified by ANCOM-BC as structural zeros (Table 3). At the genus level, 180 of 299 tested genera were FDR-significant after excluding structural-zero findings, while 43 additional genera were classified as structural zeros. Taxonomic labels for the leading ASVs are reported at genus level only where supported by the reference comparison (Table 4).

**Table 3.** Differential-abundance summary.

| Analysis | Features tested | FDR-significant non-structural-zero ASVs | Structural zeros | Higher in rampant | Lower in rampant | Notes |
| --- | --- | --- | --- | --- | --- | --- |
| Primary ANCOM-BC (prevalence $\geq 9/88$ samples) | 758 ASVs | 239 | 239 | 63 | 176 | Caries-free reference; $q < 0.05$ |
| Post hoc prevalence ($\geq 18/88$ samples) | 393 ASVs | 176 | 41 | - | - | 133/239 overlap; 100% direction concordance |
| Post hoc *S. mutans*-adjusted ANCOM-BC | - | 95 | - | - | - | 68 overlap; strong collinearity |

**Table 4.** Leading non-S. mutans ASVs.

| ASV | Genus (reference support) | ANCOM-BC coefficient (q) | Direction | Prevalence (rampant/free) | Sensitivity |
| --- | --- | --- | --- | --- | --- |
| ASV_00778 | Stomatobaculum (Strong) | -1.64 (q=2.87e-14) | Lower in rampant caries | 2.3% / 43.2% | Stricter filter: yes; adjusted model: yes |
| ASV_00351 | Lachnoanaerobaculum (Strong) | -2.33 (q=6.40e-13) | Lower in rampant caries | 2.3% / 47.7% | Stricter filter: yes; adjusted model: yes |
| ASV_00131 | Ligilactobacillus (Plausible) | 2.66 (q=4.20e-12) | Higher in rampant caries | 84.1% / 11.4% | Stricter filter: yes; adjusted model: yes |
| ASV_00136 | Porphyromonas (Strong) | -2.55 (q=5.63e-12) | Lower in rampant caries | 47.7% / 90.9% | Stricter filter: yes; adjusted model: yes |

### Post hoc analyses showed directional robustness but sensitivity of individual ASV findings

When the prevalence threshold was increased to at least 18 of 88 samples, 393 ASVs were tested, with 176 FDR-significant non-structural-zero ASVs and 41 structural-zero ASVs; 133/239 (55.65%) primary FDR-significant non-structural-zero ASVs were retained, with 100% direction concordance. In the post hoc *S. mutans*-adjusted differential-abundance model, 95 non-structural-zero ASVs were FDR-significant, including 68 overlapping unadjusted findings with 100% direction concordance. Because *S. mutans* abundance was strongly correlated with caries status, this adjusted feature-level analysis was treated as a sensitivity analysis rather than evidence of statistical independence.

## Discussion

In this public cohort, plaque community composition differed between rampant-caries and caries-free children after exclusion of one reference-supported *S. mutans* ASV. The pre-specified effect estimate was modest (pseudo-R^2^=0.0462), nearly unchanged after rarefaction, and not accompanied by evidence of unequal multivariate dispersion. Shannon diversity did not materially differ. Many ASVs met the differential-abundance criterion under the pre-specified filter, predominantly with lower bias-corrected abundance in rampant caries, but individual feature calls changed with the prevalence threshold and adjusted model.

These findings are consistent with an ecological interpretation of caries in which clinical state is associated with the configuration of plaque taxa rather than necessarily with a uniform change in a single within-sample diversity index [1-4]. Bray-Curtis composition and Shannon diversity capture different properties of a community. A group difference in composition can coexist with overlapping Shannon distributions when relative abundances differ between groups without a consistent change in richness or evenness. The present results therefore concern a community-composition difference, not a generalized decline in within-sample diversity.

The inverse association of the *S. mutans* ASV is atypical relative to the broader literature and must be interpreted cautiously. Established *S. mutans* cariogenic biology remains strongly supported by its acidogenic, acid-tolerant, and biofilm-matrix functions [2,5-7]. At the same time, community studies have reported caries with low or undetectable *S. mutans* and associations involving multiple plaque taxa [9-11]. The observed pattern is specific to this cohort and to the sequenced 16S region/reference assignment. It is not evidence against the established cariogenic role of *S. mutans*, and it provides no basis for describing *S. mutans* as protective.

Excluding the reference-supported ASV and statistically adjusting for its abundance answer different questions. The primary analysis removed that ASV from raw counts and compared the re-normalized remainder of each community. The adjusted model instead attempted to partition a group association in the presence of a covariate strongly correlated with group status. The latter association was attenuated, but the collinearity (r=-0.869; VIF=4.09) limits interpretation of a partial group term.

Neither the adjusted community result nor adjusted differential abundance establishes an association statistically independent of *S. mutans*.

The differential-abundance analysis supports a broad signal but not a fixed list of definitive taxa. Among the 758 ASVs retained by the pre-specified prevalence filter, 239 non-structural-zero ASVs were FDR-significant and 239 additional ASVs were structural zeros. Most FDR-significant non-structural-zero ASVs had lower bias-corrected abundance in rampant caries. Direction concordance among overlap in post hoc analyses was high, but the number of significant features fell with the stricter prevalence filter and after adjustment. This sensitivity is expected for sparse compositional data and means that individual ASV findings require replication rather than mechanistic interpretation.

Taxonomic resolution is another important constraint. Read 2 was selected for single-end analysis before caries-group analysis because paired reads could not overlap across the expected V3-V4 amplicon. Exact matching within the sequenced Read 2 region to eHOMD supported identification of one *S. mutans* ASV but does not make the remaining ASVs species-resolved. Accordingly, leading findings are reported at the defensible genus level, and the post hoc comparison distinguishes labels with strong versus plausible genus-level reference support. Species-level assignment would exceed the available sequence evidence.

This study has limitations. It is a secondary analysis of one public case-control cohort of 88 children, which limits precision and generalizability and cannot establish temporal ordering or causation. The original study reported matching of the case and control groups by age and sex, but individual age/sex values and matching identifiers were unavailable in the public metadata; the original matching could therefore not be incorporated into the reanalysis. Other factors potentially related to plaque ecology, including diet, oral hygiene, salivary characteristics, treatment history, socioeconomic conditions, and detailed plaque site, were not available as participant-level public inputs for this reanalysis.

Additional technical limitations deserve emphasis. The public archive supplied raw reads but not the published processed abundance table and its full derivation. The archive’s paired-spot total and the article’s total-read description are therefore not interchangeable. Amplicon counts are relative and do not establish absolute organism load; extraction, sequencing, archival, and batch effects cannot be fully evaluated. Differential-abundance results also depend on prevalence filtering and the ANCOM-BC model specification. Structural-zero ASVs and FDR-significant non-structural-zero ASVs should remain distinct.

Future work should use longitudinal, adequately powered cohorts with accessible matching metadata, plaque-site information, absolute-quantification measures, and validated species- or strain-level assays. Such studies should evaluate behavioural and environmental drivers alongside microbial patterns and distinguish prediction from explanation. Replication of the primary exclusion analysis in independent datasets would establish whether the observed community-composition difference is generalizable.

In conclusion, rampant caries in this cohort was associated with a modest plaque-community compositional difference after exclusion of a reference-supported *S. mutans* ASV, without a corresponding difference in Shannon diversity. The primary result was robust to rarefaction. The findings support a community-level ecological interpretation while retaining clear limits: individual taxa are association signals, the inverse *S. mutans* pattern is cohort-specific, and collinearity prevents a claim of statistical independence.

## Data and code availability

Raw sequencing data are publicly available through NCBI BioProject PRJNA1141721. Analysis code is available from the authors and may be deposited in a public repository in a future version.

